# A metacognitive response of the face and the heart in rats

**DOI:** 10.64898/2026.07.06.736819

**Authors:** Joana F. Doutel Figueira, Nelson K. Totah

**Affiliations:** University of Helsinki

## Abstract

Humans make emotional facial expressions and have a cardiac response when they catch themselves in a mistake or receive feedback about task performance. We tested whether rats exhibit similar visceral responses in the context of metacognition. We assessed heart rate variability (HRV) and machine learning-detected facial expressions as female and male rats detected and stopped in-progress mistakes and received post-choice rewards or error cues. HRV increased during internally detected mistakes, as well as in response to external error cues for both sexes. Errors were associated with an HRV response when parasympathetic tone was higher, while rewards were associated with an HRV response when sympathetic tone was higher. We observed sex-specific effects of cardiac interoception on cognitive control over real-time action correction, in that low parasympathetic tone was associated with reduced ability to stop in-progress mistakes exclusively in females. Rats made facial expressions during mistake detection and in response to task feedback. Outcome-related facial expressions were valence-specific, in that the facial expression after error feedback was delayed relative to the post-reward facial expression. Our results suggest that rats have a visceral experience during metacognitive monitoring.

## Introduction

Imagine catching yourself from making a mistake. You stop moving just in time. This is evidence of metacognitive control, which can be described as self-awareness of an in-progress mistake and ongoing task performance. Human subjects have a visceral response, in the form of a heart rate change, after catching themselves in a mistake^1–3^. While metacognitive monitoring of ongoing actions and task-related outcomes is a cognitive capacity of many animals, including non-human primates, rodents, and even dolphins^4–9^, there have (to our knowledge) been no reports describing whether animals have a visceral response during metacognition. Although recent work has shown that male mice make emotional facial expressions in response to passively receiving reward^10^, it is unknown if rodents produce facial expressions in response to task performance outcomes.

We developed a behavioral task for rats in which they detect and stop in-progress mistakes^11–13^ based on an analogous task in humans^14–19^. Head-fixed rats were trained to perform a Go/NoGo task on a non-motorized treadmill. In response to the NoGo stimulus, rats committed “near-mistakes” characterized by incorrectly running and then cancelling the response before crossing the response threshold (i.e., distance run). The decision to stop an in-progress incorrect response occurs without any new sensory cue. Thus, near-mistakes involve self-awareness of ongoing performance and a volitional switch between two learned responses^11–13^. We assessed visceral responses during metacognition by aligning near-mistake peak velocity with electrocardiogram (ECG) and facial videography.

## Results

We trained 4 female and 5 male rats on a Go/NoGo task while they were head-fixed on a treadmill (**Figure 1A**). Rats had to remain immobile for the inter-trial interval (0.5 to 2.0 sec) before one of two visual drifting gratings (Go stimulus or NoGo stimulus) appeared. A Go response required running past a distance threshold and a NoGo response required the rat to remain immobile for the duration of the stimulus. Hits were rewarded and False Alarms were signaled by an auditory error cue upon threshold crossing. The trial-wise velocity profiles for correct Go responses (Hits) and for incorrect Go responses (False Alarms) are shown for one example recording session (**Figure 1B**, left and middle panels, respectively). During some Correct Rejection (CR) trials, rats committed near-mistakes (NMs) in which they began running but decided to stop before crossing the distance threshold (**Figure 1B**, right panel). We observed different NM peak velocities. We divided CR trials into three sizes based on the peak velocity tercile (**Figure 1C**). For the female (or male) rats, there were 9,878 (12,233) CR trials. Out of those trials in female (or male) rats, there were 5,515 (3,981) trials in the lowest velocity tercile (T1), which corresponded to the rats sitting nearly immobile. We will refer to T1 peak velocity trials as CR trials. The two other peak velocity terciles are termed NMs. There were 2,744 (4,114) trials in the medium velocity tercile (T2) and 1,619 (4,138) trials in the high velocity tercile (T3) for females (or males, respectively). NM peak velocity was similar between females and males (Bayesian independent samples t-test, BF_10_ = 0.652 for T1, BF_10_ = 0.684 for T2, and BF_10_ = 0.512 for T3). The response trajectories were also similar, except that the males made a stronger deceleration after the NM (**Figure 1C**, indicated by arrow). During a NM, metacognition may monitor the intensity of response conflict^8,14,16,20^, error likelihood^21^, or the magnitude of volitional corrective action^12^.

**Figure 1.**
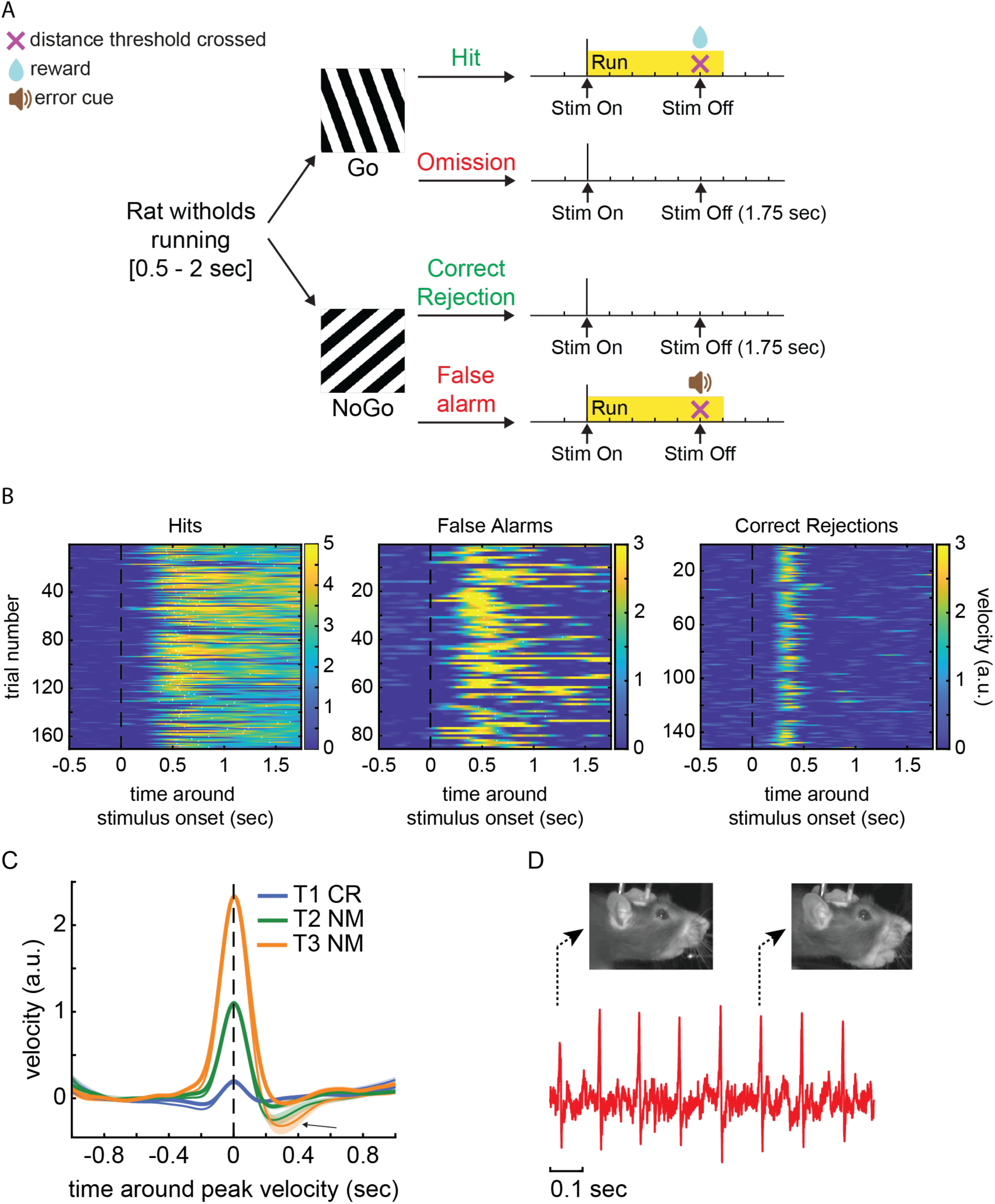
Task design and overview of data collected. **(A)** Go/Nogo task illustration. **(B)** Velocity traces aligned to stimulus onset from one session, during Hits, False Alarms and Correct Rejection trials (from left to right respectively). In the plots, each row represents one trial and, in left and middle plots, the white dots show when the distance threshold was crossed. Velocity is depicted through the colorbars, from immobility (dark blue) to fast running (bright yellow). **(C)** Average across 4 female (thick lines) and 5 male (thin lines) rats CR trials divided based on their peak velocity. Arrow on the bottom, after peak velocity, points to the deceleration that only occurs in male rats during large NMs (T3 NMs – orange thin line). The shading shows standard error. **(D)** During the task we recorded facial videography and ECG. Here we show a snippet of one session of the raw ECG and 2 timepoints of example frames of the videos collected.

We hypothesized that the cardio-visceral response during metacognition would scale with NM peak velocity. We tested this hypothesis by recording ECG and facial video (**Figure 1D**). We analyzed facial videos to detect facial expressions by performing a histogram of oriented gradients analysis for extracting image features^10^. We recorded ECG continuously throughout the entire session (**Figure 2A, B**). Given that prior work has shown a change in heart rate variability (HRV) after mistakes in humans^3^, we measured how HRV changed around the time of NM peak velocity. Out of 5 male rats, 2 had poor signal-to-noise ratio for detecting R peaks and HRV was not calculated. We calculated HRV as the root mean square of successive differences (RMSSD) between R peaks. RMSSD is robust against interference from respiration and is a direct correlate of parasympathetic nervous system control over the heart with higher RMSSD indicating increased parasympathetic nervous system activity^22–24^. We calculated RMSSD in a 500 msec sliding window (100 msec steps). This time window duration is sufficient for calculating RMSSD given that the resting heart rate of adult female and male rats is around 6 beats per second^25^. We found that average heart rate during the behavioral task (approximately 35 min duration) was 473±48.4 bpm (mean ± standard error) in females (N = 4 rats) and 463±18.5 bpm in males (N=3 rats). In females (and males), we recorded 90 (41) sessions and 5,515 (2,339) CR trials, 2,744 (2,249) T2 NM trials, and 1,619 (2,125) T3 NM trials. Average HRV in each recording session significantly varied across rats and sessions (linear mixed effects model with session count as a fixed effect and rat as a random factor, F(1, 722) = 11.33, p=0.011, **Figure 2C**). Males had higher session-averaged HRV compared to females (Bayesian independent samples t-test, BF_10_ = 54.750, **Figure 2D**). This difference may be due to female rats running more during the recording session (Bayesian independent samples t-test, BF_10_ = 4324.260, **Figure 2D**). Running, in addition to being the instrumental response in this speeded reaction time task, could also occur during the inter-trial interval (ITI) during which rats were allowed to run freely (with the penalty of delaying the stimulus onset until the rat remained immobile for at least 500 msec).

**Figure 2.**
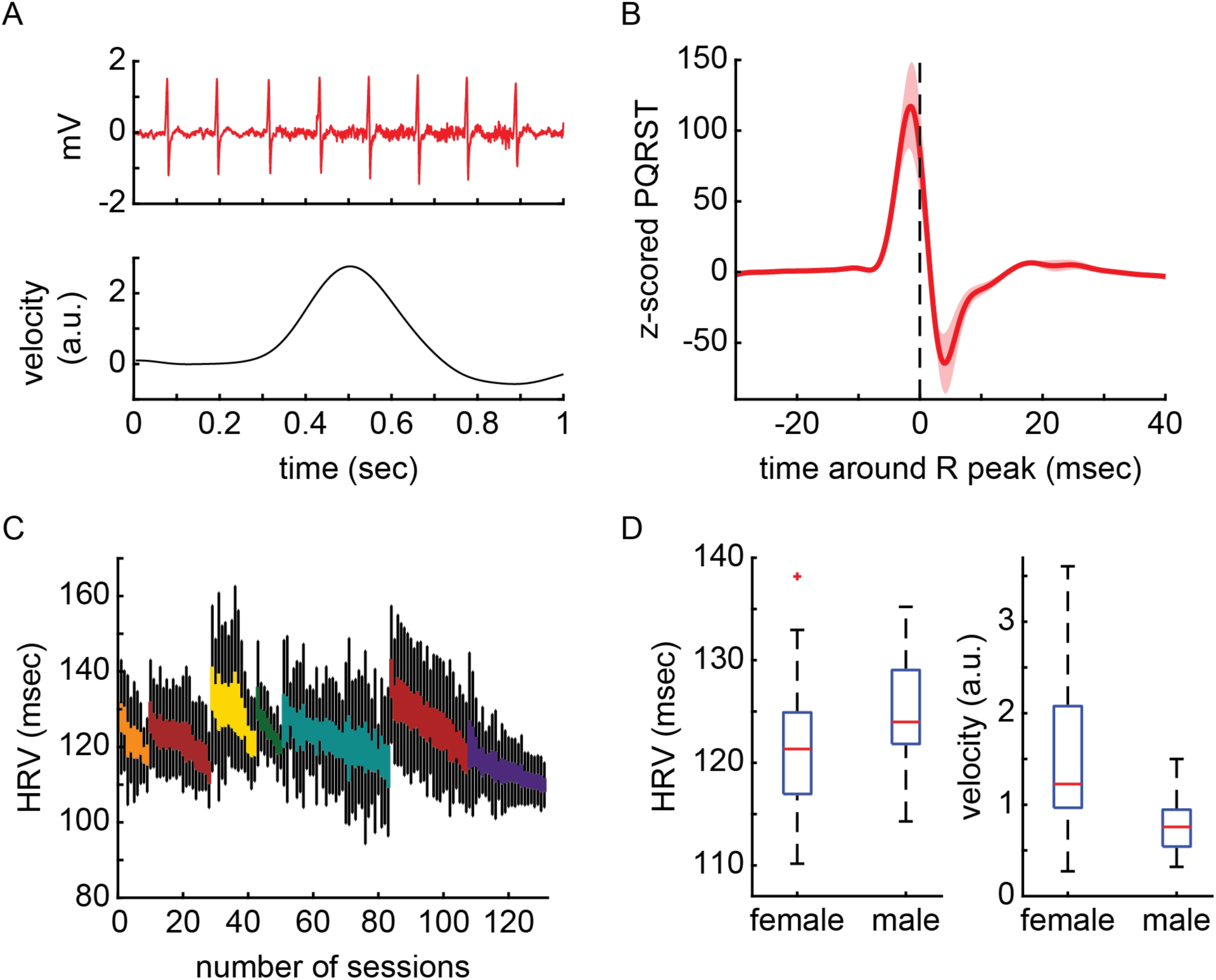
HRV dynamics of female and male rats during the Go/Nogo task. **(A)** 1-second example of ECG aligned to velocity during one session. **(B)** Average of ECG data aligned to the R peak of the QRST wave. The shading shows standard error across 7 rats. **(C)** Averaged HRV of each session. Colors represent the different rats. **(D)** Averaged HRV and velocity across female (N=4) and male (N=3) rats.

### HRV is modulated during near-mistakes and after task outcomes

We assessed HRV dynamics around the NM peak velocity. Given that HRV values were session-specific (**Figure 2C**), we present HRV z-scored to the mean and standard deviation of the recording session. HRV changed during NMs for both sexes (**Figure 3A, B**). There was a peak velocity tercile X time interaction (2-way Bayesian repeated-measures ANOVA, BF_10_ = 1.247x10^33^ for females and BF_10_ = 6.935x10^48^ for males). As the rat prepares to run on T3 NM trials, HRV increases, and then decreases during running before again increasing as the rat slows down (**Figure 3C, D**).

**Figure 3.**
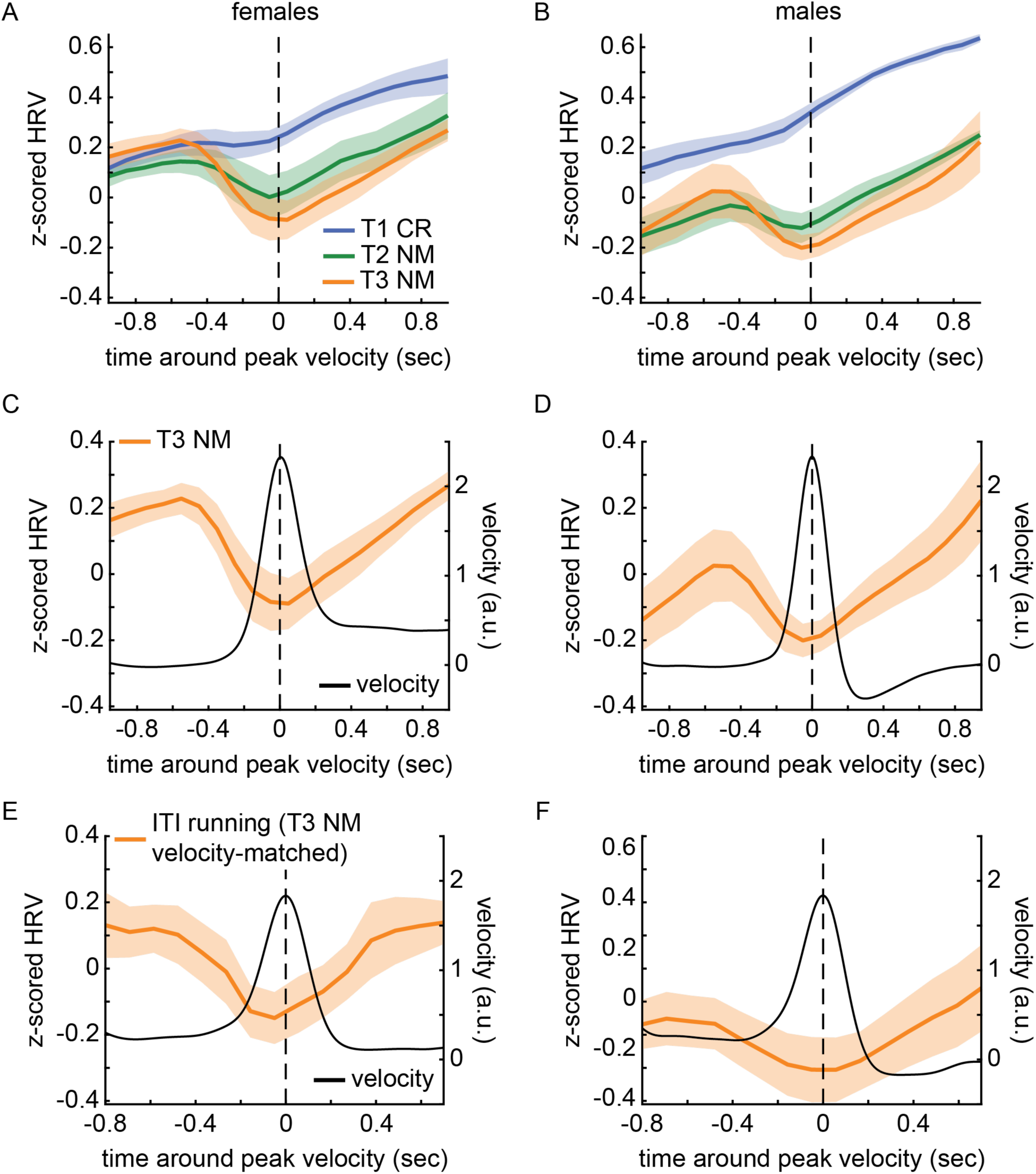
HRV is modulated during NMs in female and male rats. Panels (A) and (B) show z-scored HRV aligned to peak velocity in female and male rats, respectively. Each line represents a tertile of CR based on their peak velocity. The shading illustrates standard error across 4 female (A) and 3 male (B) rats. In panels (C) and (D), z-scored HRV during T3 NMs is shown together with velocity for female and male rats, respectively. Orange line shows z-scored HRV while velocity is shown through the black line. The shading on the orange line shows standard error across 4 female (C) and 3 male (D) rats. Panels (E) and (F) depict the z-scored HRV (orange line) and velocity (black line) during ITI running aligned to times matching T3 NM peak velocity for female and male rats, respectively. The orange shading shows standard error across 4 female (E) and 3 male (F) rats.

We tested whether the observed HRV dynamics during NMs is related to physical activity, rather than metacognitive monitoring of an in-progress mistake. To do so, we analyzed HRV around bouts of running during the ITI that were velocity-matched to NMs (**Figure 3E, F**). Although ITI T3 running bouts were also associated with HRV changes over time, this differed from NM T3 running bouts (2-way Bayesian repeated-measures ANOVA, BF_10_ = 5.608x10^1^^4^ for females and BF_10_ = 2.566x10^1^^3^ for males). Thus, HRV changes during running, but it changes more so when that running is associated with a NM. We conclude that metacognition independently effects HRV in addition to running.

We also characterized HRV dynamics during another aspect of metacognitive awareness of task performance: the receipt of task feedback (i.e., error cues and rewards). Across 4 female (or 3 male) rats, there were 8,032 (4,351) error cues and 15,894 (9,913) rewards over 90 (41) sessions. In False Alarm trials, the rats ran past the distance threshold (triggering an audible error cue), and then quickly slowed down, whereas in Hit trials, the rats gradually slowed down while consuming reward (**Figure 4A, B**, lower panels). The rats were given a 2 sec task time-out after a False Alarm and additional running or immobility would not alter trial outcome. Therefore, the rapid stopping after the error cue may be a sign of metacognitive awareness.

**Figure 4.**
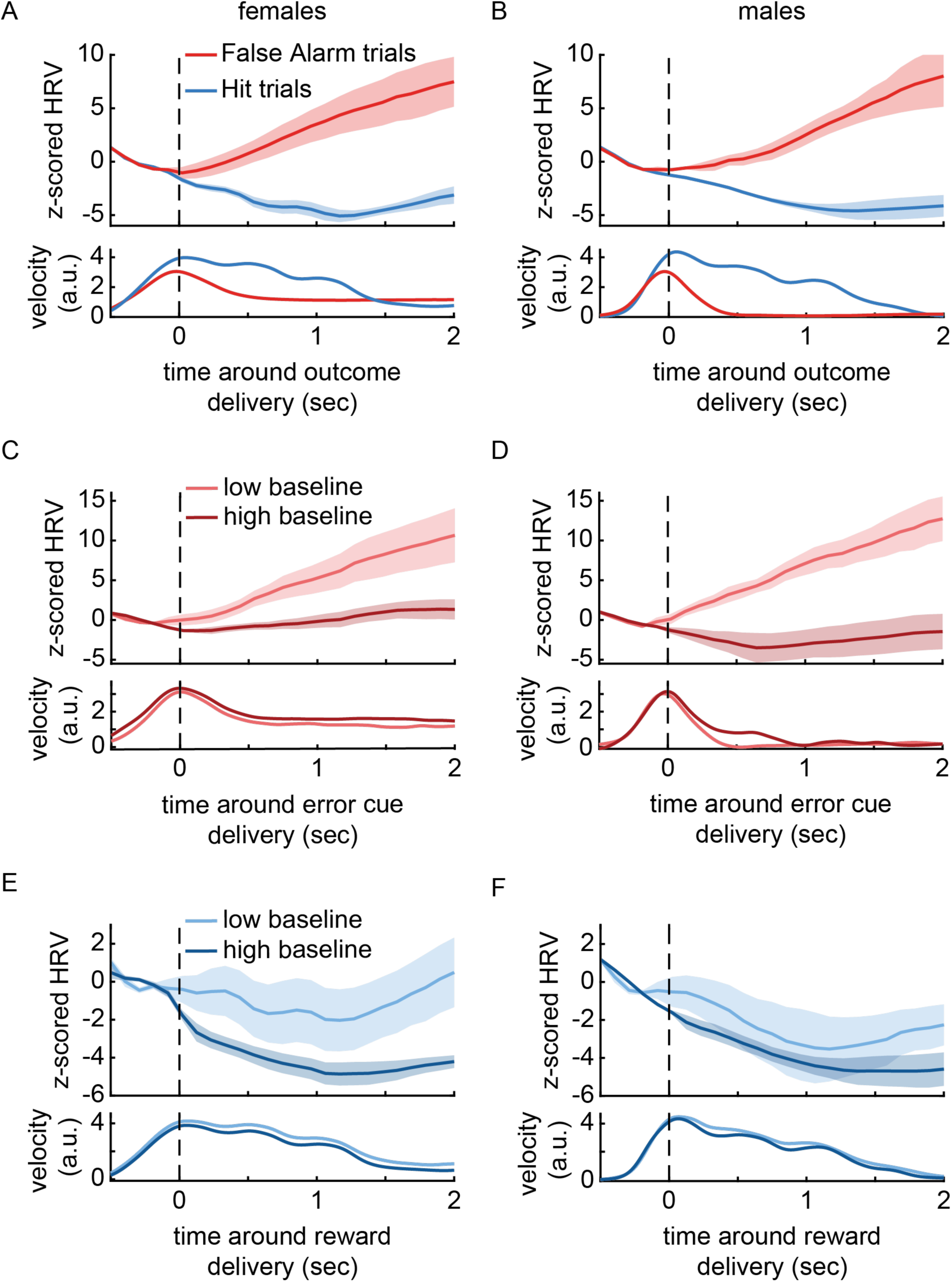
During task performance outcomes HRV is metacognitively modulated in female and male rats and depends on pre-stimulus HRV. The lines in the upper plots show z-scored HRV and lines in the lower plots show average velocity, for female **(A)** and male **(B)** rats. Blue and red lines are, respectively, Hit and False Alarm trials aligned to outcome delivery. In the upper plots shading show standard error across 4 female **(A)** and 3 male **(B)** rats. . False Alarm trials were divided based on their pre-stimulus HRV into high (top 25%) and low (bottom 25%) baseline for female **(C)** and male **(D)** rats. Upper and lower panels show z-scored HRV and velocity, respectively, aligned to outcome delivery. The shading on the upper plot shows standard error across 4 female **(C)** and 3 male **(D)** rats. Panels **(E)** and **(F)** show Hit trials divided based on their pre-stimulus HRV into high (top 25%) and low (bottom 25%) baseline for female and male rats, respectively. Upper panels show z-scored HRV aligned to outcome delivery. The shading represents the standard error across 4 female **(E)** and 3 male **(F)** rats. Lower plots show the respective velocity aligned to outcome delivery.

In both sexes, the HRV increased after the error cue, whereas on Hit trials, HRV decreased (**Figure 4A, B**, upper panels). Although both Hits and False Alarms are associated with slowing after the outcome cue, the HRV diverges in opposite directions. This is further evidence that HRV changes, while affected by running, are also affected by cognitive context. Overall, we conclude that, although there are running-related HRV changes, there is a metacognitive modulation of HRV riding on top of running-related changes that are associated with monitoring of in-progress mistakes and task performance outcomes.

### Metacognition-related HRV modulation depends on pre-stimulus balance between the parasympathetic and sympathetic nervous system

Changes in HRV that are evoked by task events are dependent on the ongoing state of the parasympathetic nervous system in humans^26^. Therefore, we tested if the ongoing state of the parasympathetic nervous system at stimulus onset influences the HRV in response to task feedback and during NMs. We defined states of low or high parasympathetic nervous system activity by dividing trials into the bottom 25% and top 25% of pre-stimulus HRV values. For each trial, the value was the average HRV in the 500 msec prior to stimulus onset. We used all trials (Go and NoGo stimuli and independent of the response) to capture the session-wide variance in parasympathetic nervous system state. We found that the HRV increase after error cues was observed in both sexes when baseline HRV was low (i.e., greater relative activation of the sympathetic nervous system relative to the parasympathetic nervous system) (**Figure 4C, D**). On the other hand, the decrease in HRV after rewards was primarily observed when baseline HRV was high (i.e., heightened parasympathetic relative to sympathetic nervous system tone) (**Figure 4E, F**). Our results show that the HRV increase versus decrease after error cues or rewards, respectively, was differentially dependent upon the pre-stimulus parasympathetic nervous system state. We also characterized the relationship between ongoing parasympathetic nervous system state and the HRV dynamics during NMs. We assessed only T3 trials because, in those trials, we saw the largest HRV dynamics (**Figure 3C, 3D**). We found that a higher pre-stimulus HRV was associated with more dynamic HRV pattern during the NM in both females and males (**Figure 5A, B**). Thus, metacognition-related HRV changes are influenced by the ongoing parasympathetic nervous system state.

**Figure 5.**
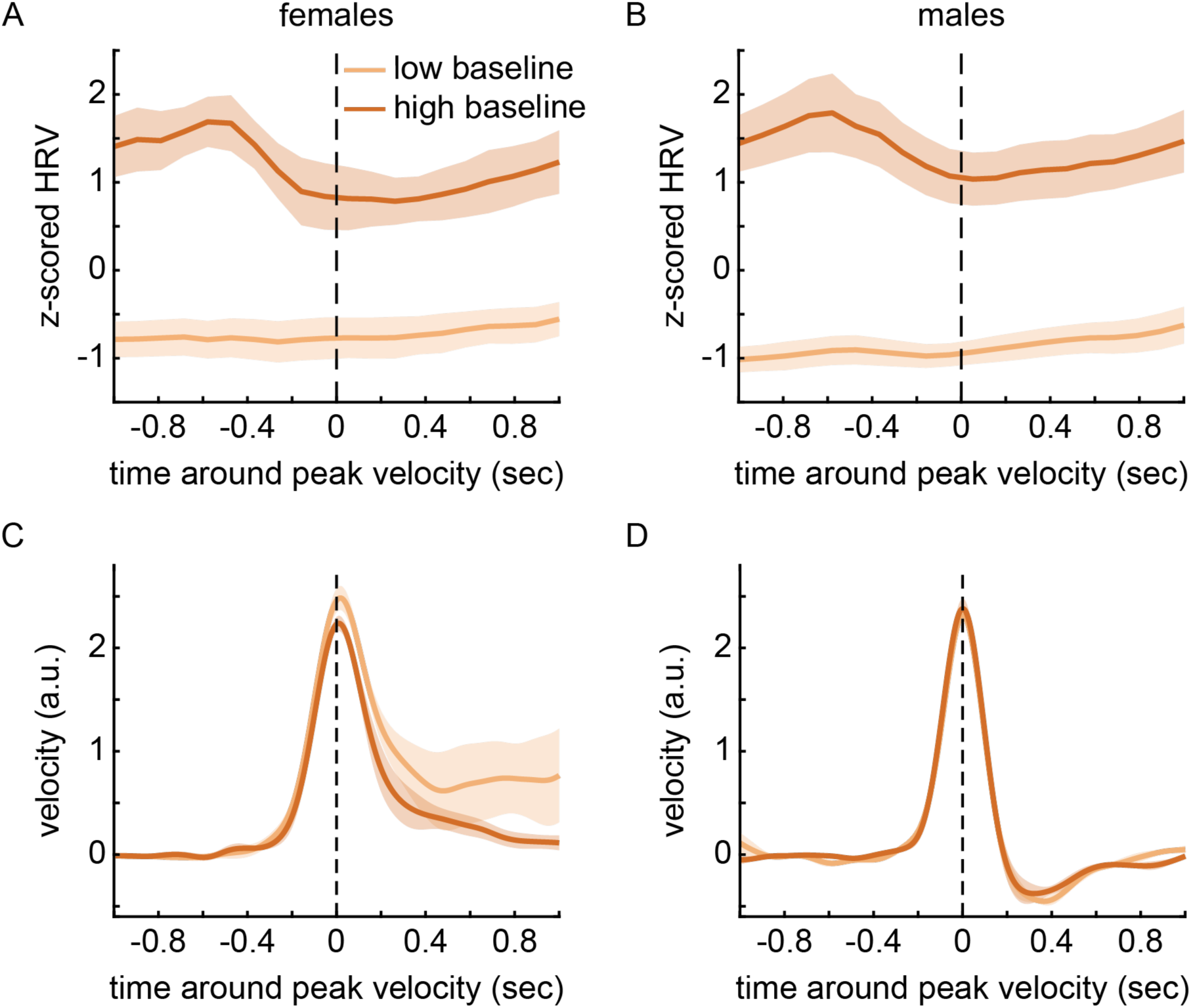
Pre-stimulus HRV influences the metacognition-related HRV modulation during T3 NMs in female and male rats, and it affects the behavior itself but only in females. T3 NMs trials were divided based on their pre-stimulus HRV into high (top 25%) and low (bottom 25%) baseline for female and male rats. Z-score HRV aligned to peak velocity is shown in panels **(A)** and **(B)**, for female and male rats, respectively. The shading in both panels show standard error across female (N=4) and male (N=3) rats. Panels **(C)** and **(D)** show the velocity for low and high baseline T3 NM trials aligned to peak velocity.

### Cardiac interoception affects metacognitive behavior in only female rats

The analyses presented thus far demonstrate that metacognition is associated with a cardiac response in rats. However, it is also possible that cardiac activity state may modulate performance monitoring and behavior (i.e., cardiac interoceptive effects on cognition). We noticed that the NM velocity appeared to depend on pre-stimulus HRV, specifically, in female rats. When ongoing HRV was low, female rats made larger NMs (Bayesian paired samples t-test on T3 NM peak velocity, BF_10_ = 5.105) and they did not fully return to immobility within the 1 sec following NM peak velocity (**Figure 5C**). In other words, in a heightened sympathetic nervous system (fight-or-flight activation) state, female rats came closer to making mistakes and, although they managed to slow down and prevent the mistake, they had difficulty returning to immobility. On the other hand, males had similar peak velocity (Bayesian paired samples t-test on T3 NM peak velocity, BF_10_ = 0.842, indicating anecdotal evidence in favor of the null hypothesis) and trajectory after peak velocity regardless of the pre-stimulus HRV (**Figure 5D**). Thus, dampened parasympathetic nervous system (and heightened sympathetic nervous system state) favoring a heightened arousal modulates metacognition-related behavior in female rats.

### Rats make a facial expression during metacognitive awareness of errors

Recent work has shown that mice make “emotional” facial expressions when passively receiving stimuli that would evoke emotional facial expressions in humans, such as sucrose reward, fear cues, and pain^27,28^. Given that metacognitive awareness is associated with emotional facial expressions in humans^29^, we assessed whether rats make facial expressions during metacognitive monitoring of NMs, rewards, and error cues. We tested the hypothesis that features of the facial image during NMs would become dissimilar from the facial features observed during CR trials in which no incorrect running occurred and, thus, metacognitive monitoring of errors is absent. Across 4 female (or 5 male) rats, there were 5,169 (3,979) T1 trials, 2,534 (4,109) T2 trials, and 1,491 (4,138) T3 trials over 83 (74) sessions.

We described the main features of the facial image using a histogram of oriented gradients (HOG) analysis of each image frame (**Supplementary Figure 1**). Once each frame was represented as a HOG, we compared each frame in a -0.8 to +0.7 sec window around T2 or T3 NM peak velocity to the average HOG during CR trials. This CR face was calculated by averaging the HOGs across frames within each trial (from NoGo stimulus onset until the end of the trial, 1.75 sec later) and then averaging those HOGs across trials within each session. The dissimilarity between the facial features around NM peak velocity and the CR face was calculated as a pixel-by-pixel Pearson’s correlation coefficient (see Methods section for detailed description and **Supplementary Figure 2**). Thus, for each frame during the NM, there is a correlation value describing how similar that frame was to the average facial features when rats saw the NoGo stimulus and remained nearly immobile.

We observed a decrease in the z-score normalized correlation coefficient on T2 and T3 NMs around the peak velocity in both females (**Figure 6A**) and males (**Figure 6B**). The decrease in correlation scales with the size of the NM. A Bayesian repeated measures 2-way ANOVA indicated that our data are extremely strong evidence for a difference between NM sizes over time (time X NM size interaction: BF_10_ = 2.748x10^77^ for females and BF_10_ = 5.987x10^187^ for males). We also compared HOGs for T2 and T3 NMs and found that the T3 facial expression becomes dissimilar from the T2 facial expression (**Figure 6C, D**) These results show that rats make a facial expression during metacognitive monitoring of in-progress mistakes and that distinct facial expressions are made for different intensities of mistakes.

**Figure 6.**
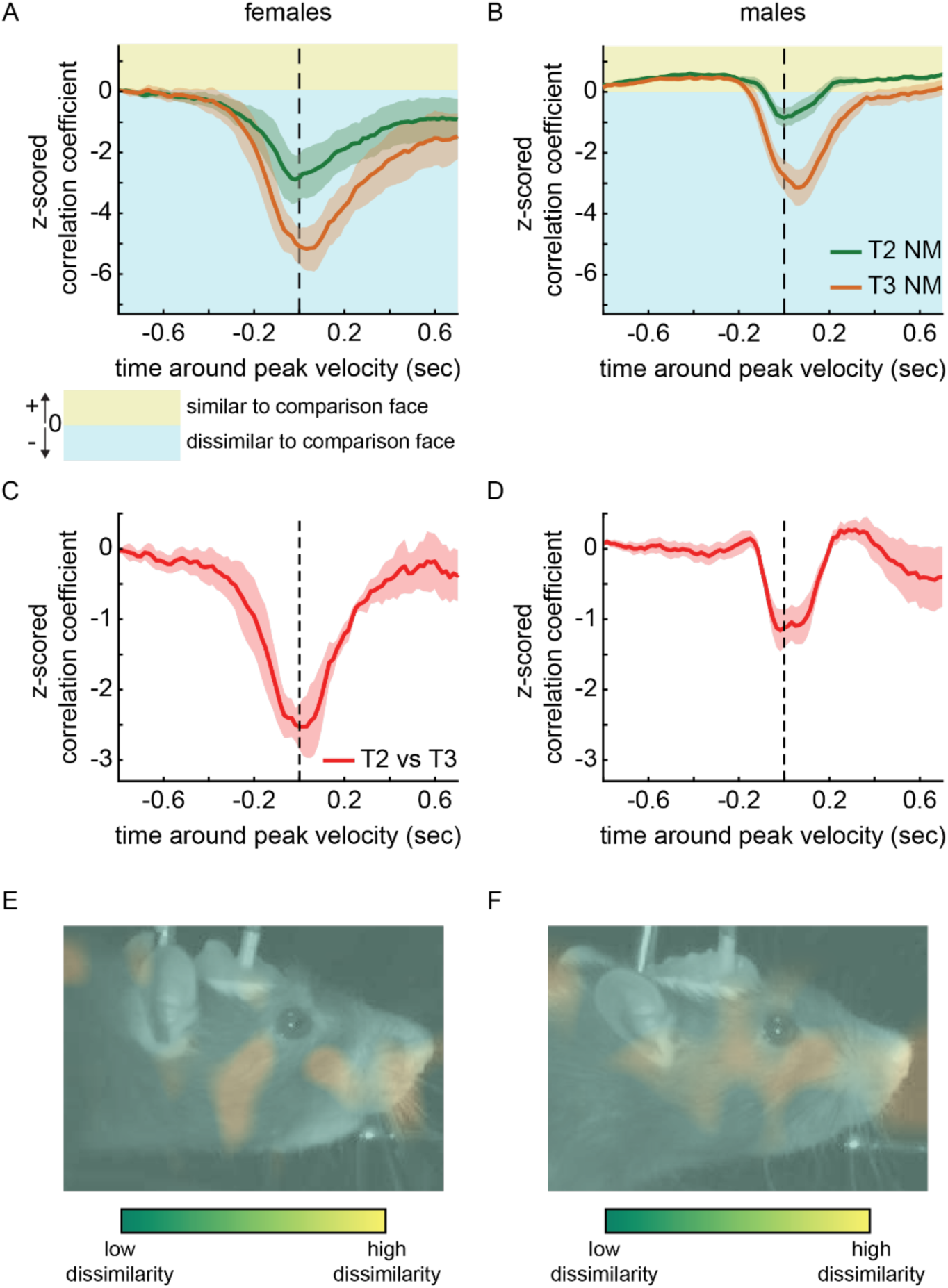
Rats make a facial expression during NMs. T2 and T3 NMs trials HOGs were correlated with an averaged HOG during T1 CR trials. Panels (A) and (B) show the z-scored correlation coefficient aligned to peak velocity for female and male rats, respectively. The plot lines shading shows standard error across 4 female (A) and 5 male (B) rats. The blue and yellow shading across the plots illustrate whether the facial expression during T2 and T3 NMs was similar (yellow) or dissimilar (blue) to the T1 CR face. Panels (C) and (D) represent where the face during peak velocity on T3 NMs (-0.05 sec to 0.1sec around peak velocity time) is most dissimilar with the T1 CR face for both female and male rats, respectively. The frames are example frames on T3 NMs. On the heatmap, green and yellow represent low and high dissimilarity, respectively.

We determined the facial features that contributed the most to the facial expression during T3 NMs by plotting a heatmap illustrating which areas of the facial image had the greatest dissimilarity compared to the CR face (**Figure 6E, F**). In females (**Figure 6E**), the T3 NM-related facial expression involves areas around the nose, cheek and ears. In males (**Figure 6F**), the highest level of dissimilarity are also around the nose, and ears. Both female and male rats show changes in similar facial regions during metacognitive detection of error.

It is possible that this dissimilarity of the face during NMs, in comparison to the immobility on CR trials, is observed because physical activity affects facial features. We tested for this possibility by comparing bouts of running during the ITI (velocity-matched to NM peak velocities) with epochs of immobility during the ITI. For both sexes, we observed a decrease in face similarity around peak velocity for T2 and T3 ITI running bouts in comparison to immobility during the ITI (**Figure 7A**, **B**). Thus, the facial expression made during NMs appears to be one that is made in general during physical activity. On the other hand, it is also possible that a metacognition-related facial expression could co-occur with a change in facial features due to physical activity. We tested this by calculating the correlation coefficient between NMs and the average HOG during velocity-matched ITI running bouts. If the facial expression is a change in facial features that occurs, in general during running, then the correlation coefficient should be positive when comparing velocity-matched bouts of running during the ITI and NMs. However, the correlation became negative immediately prior to peak velocity (**Figure 7C, D**). A Bayesian one-sample, one-sided t-test against a test value of 0 indicated strong evidence for dissimilarity of the facial expression in both sexes on T3 trials (females BF_-0_ = 7.842, males BF_-0_ = 41.41). During this epoch, the running dynamics are matched, but cognitive context differs: for NM running bouts, the rats are detecting an in-progress mistake and switching between two learned, stimulus-guided responses. The negative correlation around peak velocity suggests that metacognitive error awareness is associated with a facial expression distinct from a chance in facial features due to physical activity.

**Figure 7.**
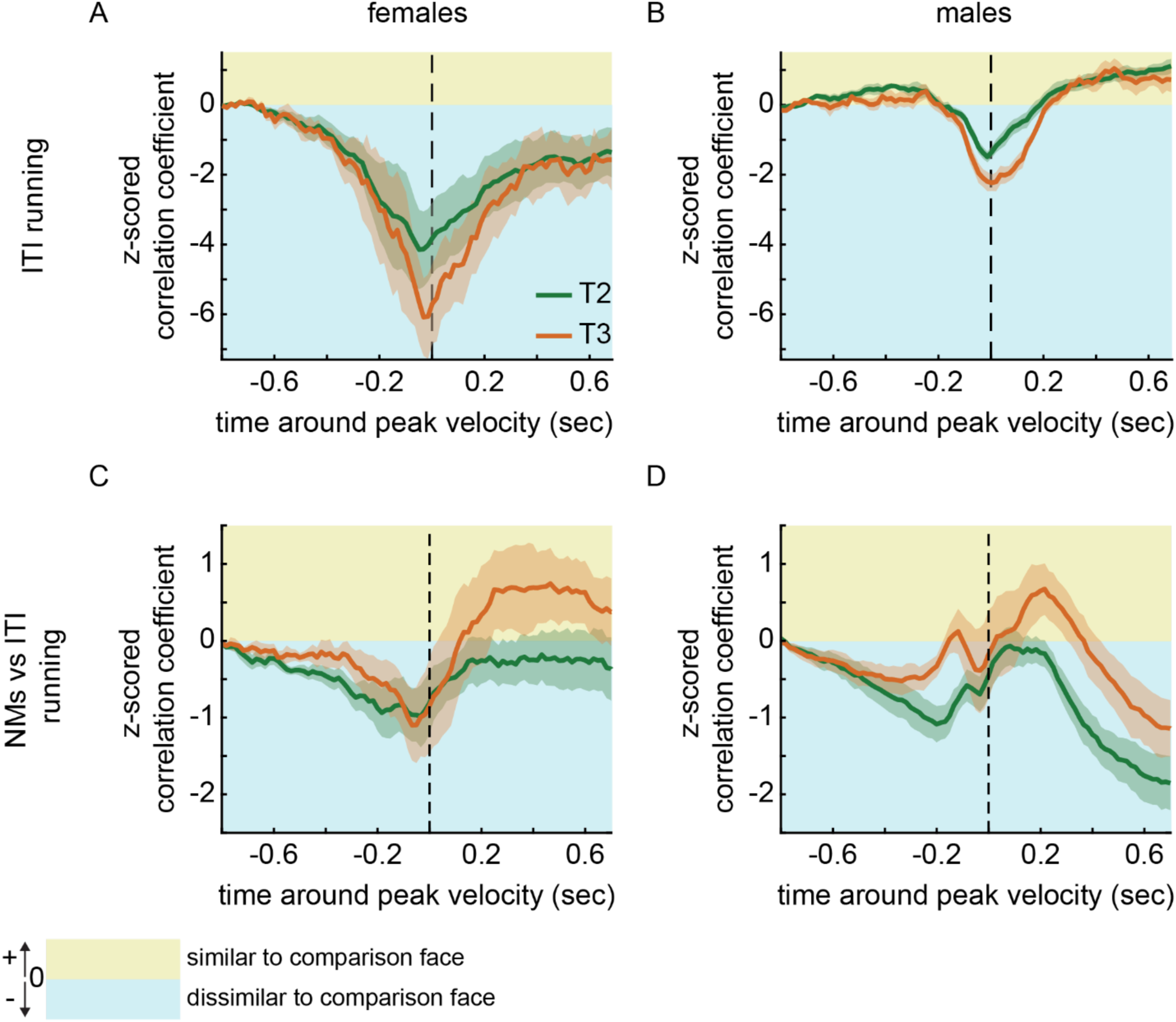
Facial expression during NMs is different than facial expression during velocity-matched running bouts, indicating that there is a metacognitive-related facial expression during NMs. Panels **(A)** and **(B)** depict the z-scored correlation coefficient between ITI running aligned to times matching T3 (or T2) NMs and T1 CR trials for female and male rats, respectively. The plot lines shading shows standard error across 4 female **(A)** and 5 male **(B)** rats. The blue and yellow shading across the plots illustrate whether the facial expression during ITI running aligned to velocity-matched T2 and T3 peak velocities was similar (yellow) or dissimilar (blue) to the ITI running aligned to velocity-matched T1 peak velocity. In panels **(C)** and **(D)** T3 (or T2) NM trials are correlated with ITI running aligned to velocity-matched T3 (or T2) for female and male rats, respectively. The plot lines shading show standard error across 4 female **(C)** and 5 male **(D)** rats. The blue and yellow shading across the plots illustrate whether the facial expression during T3 (or T2) NMs was similar (yellow) or dissimilar (blue) to the ITI running aligned to velocity-matched T3 (or T2) peak velocity.

### Facial reactions to errors is delayed and initially confused with reward

Male mice have been shown to make different facial expressions to stimuli with positive (i.e., rewards) versus negative valence (i.e., fear cues and noxious stimuli)^10^. This prior work delivered these stimuli passively without conditioning stimuli upon task performance. We looked for facial expressions related to outcomes of task performance (i.e., rewards and error cues) in female and male rats. We generated a prototypical reward outcome face and a prototypical error outcome face. We then calculated the similarity (i.e., Pearson’s correlation coefficient) of the HOGs between these prototypical faces and trial-by-trial rewards or error cues. The prototypical faces were generated by averaging the HOGs in the 2 sec after outcome delivery on a single trial and single rat (sex-specific). These procedures for generating the prototypical faces were identical to the method used in prior work in mice to promote cross-species, cross-laboratory comparison of results^10^. We found that the choice of rat and trial (e.g., first, middle, or last in the session) for generating the prototype face can change the magnitude of the correlation coefficient (**Supplementary Figure 3 and 4**), but in most cases the results are similar. For each outcome event, we calculated the frame-by-frame HOG similarity to each prototype HOG in a -0.5 to 2.0 sec window around the event. For the 4 female (or 5 male) rats, there were 7,397 (7,797) error cues and 14,657 (18,334) rewards across 83 (74) sessions. The correlation coefficients around event onset were trial-averaged within session and then averaged across sessions within rat.

Prior work in male mice shows different facial expressions to passively delivered positive and negative stimuli^10^. We tested whether the same occurred in a different species (rats), and in the context of task performance as opposed to passively delivered stimuli. When reward was delivered (**Figure 8A, B**), the average facial expression across female and male rats became more similar to the prototypical reward face (i.e., a relatively higher correlation coefficient) as the time of reward approached. The correlation remained elevated for approximately 1.5 sec. This sustained facial expression may be due to the task design: three rewards were delivered every 500 msec after distance threshold crossing on Hit trials. On these same rewarded trials, the similarity to the prototypical error cue face decreased continuously for approximately 1.5 sec. These results suggest that, around reward deliveries, female and male rats make a reward-specific facial expression that is not similar to the prototypical facial expression made after error cues. Unexpectedly, as the time of the error cue approached (**Figure 8C, D**), facial expressions became more similar to the prototypical reward face and less similar to the prototypical error cue face. However, soon thereafter, this pattern inverted: correlations to the prototypical reward facial expression became negative (dissimilar) while becoming positive (similar) to the prototypical error cue facial expression. Therefore, female and male rats initially make a reward-related facial expression regardless of whether the outcome of the trial is an error cue or a reward, and only later do they express a valence-specific facial expression.

**Figure 8.**
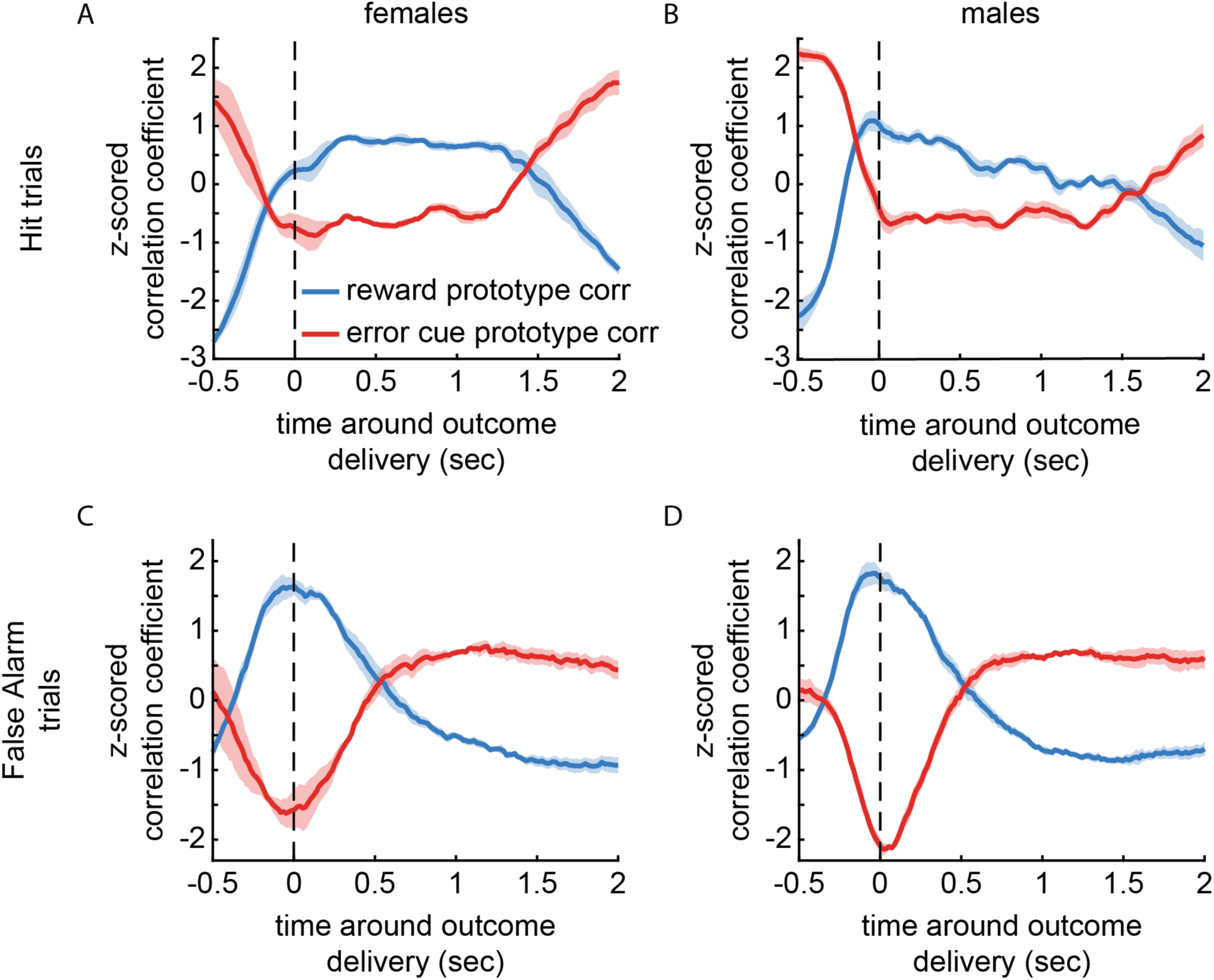
Rats make a facial expression during task performance outcomes and during errors, facial expression is delayed and initially confused with reward. False alarm trials are plotted in panels **(A)** and **(B)** for females and males, respectively. Red lines show the correlation between the facial expressions during False Alarm trials and the error cue prototype facial expression, while blue lines show the correlation with the reward prototype facial expression. The shading shows standard error across 4 female **(A)** and 5 male **(B)** rats. Facial expressions correlation between Hit trials and prototype faces are plotted in panels **(C)** and **(D)**. Red and blue lines depict the correlation with error cue and reward prototype facial expressions, respectively. The shading shows standard error across 4 female **(C)** and 5 male **(D)** rats.

## Discussion

Here, we tested whether rats, like humans^1–3^, make facial expressions and experience a cardiac HRV response during metacognitive monitoring of task performance. Metacognition occurred in two contexts: (*i*) the detection and on-line correction of an in-progress incorrect response and (*ii*) the detection of task outcomes (rewards and error cues). We found that HRV was modulated during NMs in female and male rats. The modulation scaled with the peak velocity of the NM. Critically, this HRV response was not as large when rats stopped bouts of running during the ITI, suggesting that the response was not related only to physical activity. Therefore, HRV changes in rats as they detect and stop in-progress mistakes. Recent work in human subjects has shown that proprioceptive errors are also associated with an autonomic nervous system response, including an HRV change that scales with the magnitude of the error movement^30^. While the rats were performing a decision-making task and the human participants were only experiencing proprioceptive errors, similar proprioceptive processing may play a role in how rats assess error likelihood on the treadmill. During reward delivery, HRV decreases both in female and male rats, suggesting an increase in arousal, which is consistent with prior human work where momentary positive affect was correlated with a lower HRV^31^. On the other hand, after error cues, HRV increased. Thus, our results show that metacognition is associated with a response of the heart in rats.

The ongoing state of the autonomic nervous system influenced the HRV response during metacognitive awareness of task performance in opposite directions depending on valence. For internal detection and correction of in-progress errors, as well as in response to external error cues, higher parasympathetic nervous system activity was associated with greater HRV changes. In contrast, the HRV response to rewards was larger when the sympathetic nervous system was more active. In sum, negative valence events (near-mistakes and error cues) were associated with an HRV response when rats were in a parasympathetic (less aroused, or relaxed) state, whereas positive valence events (reward delivery and consumption) were associated with an HRV response when rats were in a sympathetic (flight-or-flight) state.

We observed a sex-specific effect of cardiac interoception on cognitive control over real-time action correction. We found that female rats with a low pre-stimulus HRV made larger NMs and inhibited their mistaken response more slowly. This suggests that cardiac interoception modulates NM response dynamics, but only in female rats. However, without neural recordings and causal neural manipulations, we cannot exclude that a brain region (or circuit) provides shared drive over autonomic nervous system state and behavior rather than interoception of the heart affecting subsequent behavior.

In addition to affecting the heart, metacognition in rats was also associated with facial expressions. At the peak velocity of NMs – just as the rats were realizing that they were committing a mistake – the face became dissimilar from the face made during correct rejection trials. The facial expressions were distinct for different intensities of near-mistakes. Critically, this facial expression was also dissimilar from the face at the same time point during velocity-matched bouts of running during the ITI. These results, observed in both female and male rats, indicate that real-time monitoring of mistakes is associated with distinct facial expressions.

Female and male rats made valence-specific facial expressions in response to task outcomes. When receiving reward, rats made a facial expression that was different from the facial expression made in response to error cues. On the other hand, after an error cue, rats initially made a facial expression resembling reward delivery, followed by an error cue-specific facial expression. Therefore, we show a valence-specific facial response during metacognitive monitoring of task outcomes. These findings are in line with prior work in male mice showing valence-specific facial expressions during passive exposure to positive and negative events^10^. To our knowledge, prior rodent studies have not characterized facial expressions in response to task performance outcomes, and instead used facial expressions to predict reaction times and decisions^32,33^.

The behavioral task used in this study permitted, in rats, assessment of metacognitive monitoring of task performance. We assessed two types of metacognitive monitoring: real-time detection of mistakes and detection of task outcomes. Humans and non-human primates have been shown to monitor, detect, and perform real-time correction of mistakes^14–19,34–37^, and we now show this ability in rats. These species can also assess decision confidence and to make choices based on task context and their ongoing task performance^4,6–8^. For instance, when performing a difficult discrimination between sensory stimuli, humans, rodents, and even dolphins appear to understand that they might make an error and will take advantage of an opportunity to skip that decision and instead perform an easier discrimination^5,7,9^. Our task cannot distinguish whether metacognitive awareness during NMs is related to response conflict monitoring^20^ or awareness of error likelihood^21^, but brain regions involved in monitoring conflict and error likelihood such as the anterior cingulate cortex and the insula cortex may be involved in modulating the autonomic nervous system and in-turn affecting HRV and facial expressions.

How we behave and think affects how we feel in a visceral sense. When we catch ourselves in a mistake or become aware of the consequences of our decisions, we make facial expressions and might feel our heart flutter (accompanied by an HRV change). Yet it is not just humans that have bodies -- every multi-cellular organism has some form of physical organization resembling a body. Therefore, the mind-body connection could be present in other species. Our results show that this is true for rats, which have a response of the face and of the heart during metacognitive awareness of errors and task outcomes.

## Material and Methods

### Experimental model

4 female and 5 male Lister-Hooded rats (140 – 196 g body weight when implanted with cardiothoracic electrodes and a head chamber with head-fixation post). The rats were supplied by either Charles River or Envigo (Inotiv). Rats were pair housed and handled daily for 5 to 10 minutes for 5 to 14 days prior to surgery. After surgery, rats were single housed in a reversed light-dark cycle (08:00 lights off, 20:00 lights on). Behavioral training and task performance occurred during the rat’s active phase. All procedures were carried out after approval by local authorities (license number: ESAVI/6859/2024) and following the European Community Guidelines for the care and Use of laboratory Animals (EU Directive 2010/63/EU).

### Surgery

Anesthesia was induced with a i.p. injection of ketamine (80 mg/kg) and medetomidine (0.01 mg/kg) and maintained with isoflurane (∼0.1 – 1.0% in O2). We prefer to use this combination for induction of anesthesia for three reasons. First, it reduces the risk of isoflurane overdose. Second, it reduces the stress experienced by the animal (the treatment is a better alternative than placing the animal in an induction box with an unpleasant-smelling gas that causes loss of consciousness). Third, compared to isoflurane induction, this combination allows more time to position the rat on the operating table and attach the stereotactic frame and vital signs monitor. Heart rate and SpO2 were monitored (Somnosuite, Kent Scientific). A half dose of ketamine and medetomidine was administered as needed during surgery if the level of anesthesia was too superficial. Ketamine alone (0.1 mL of a 100 mg/ml solution) was injected i.m. if the anesthesia became unstable and difficult to control with isoflurane; or if isoflurane flow was already high (e.g., >1.0%) but respiratory rate and heart rate were increasing; or if the heart rate was stable but the rat reacted to painful stimuli (taking advantage of the analgesic effect of ketamine). Buprenorphine (0.05 mg/kg, s.c.) and carprofen (5.0 mg/kg, s.c.) were administered for analgesia. Dexamethasone (2.0 mg/kg, s.c.) was administered to reduce respiratory inflammation.

After establishing a stable, deep anesthetized state and administering analgesics, the head was shaved and rubbed with betadine. Lidocaine (10 mg/kg) and adrenaline (10 ug/kg) were injected s.c. over the skull. We made a small mediolateral incision around 1 cm rostral to the posterior edge of the skull taking care to not expose neck muscle. Connective tissue was removed by lifting and cutting away tissue. Absorbent gelatine sponge was inserted into the wound (Ethicon, Spongostan Film, MS001). The rat was then placed into a supine position and the chest was shaved and rubbed with betadine. An approximately 1 cm long incision was made to the anterior chest to the right of the sternum. The skin was lifted with one forceps (Fine Science Tools, 11210-20) and then another forceps (Fine Science Tools, 11254-20) was inserted into the incision and opened to break connective tissue and fat at the opening of the incision. A feeding cannula (Fine Science Tools, 16 gauge, 18061-75) was inserted through this opening and gently pushed subcutaneously towards the open skin over the head. The cannula was run behind the right ear. The rat was then laid on its side and the EKG wire (A-M Systems, 793200) was threaded through the cannula starting from the incision over the head. The other end of the wire had been already attached to an electrode interface board (Neuralynx, EIB-8). Once the wire was through the cannula, the cannula was retracted out of the thoracic incision while leaving the wire in place. A small ball of glue (Loctite 401 glue) was put on the tip of the wire taking care to not cover the entire uninsulated portion of wire. The conjunctive tissue was pulled away using forceps (Fine Science Tools, 11254-20) to expose the underlying muscle. The wire was guided into an orientation parallel to the muscle fibers and then sutured onto the muscle (Seralene 6/0 DRT-12, reference: LO07340B) immediately anterior to the ball of glue. The skin was sutured (Ethilon 5-0 FS-3, reference: 1628H). This procedure was then repeated at the posterior chest to the left of the sternum. The wire inserted there can serve as a reference for differential recording (if needed). Once both wires were in place and the skin was sutured, the rat was turned back into prone position to implant the head chamber and head-fixation post.

Lidocaine (s.c.) was injected in the skin overlaying the skull. The incision was opened with forceps and scissors (Fine Science Tools, 14060-09) were used to cut the skin in the anterior direction running along the left and right edges of the skull. Care was taken to not expose the temporal muscles. The skin was cut until immediately posterior to the eyes. A cut connecting the left and right lateral incisions was made in the mediolateral direction to remove the skin flap. The skin and connective tissue were removed to expose the skull. The wound margin was cauterized and the exposed bone was wiped dry and cleaned with 5% hydrogen peroxide. The hydrogen peroxide was washed away with copious amounts of sterile saline (e.g., 20-40 mL). The bone surface was scratched using a scalpel (Swann-Morton, 0503) in a grid pattern to facilitate adhesion for the UV light polymerizing cement used to affix the implant to the skull. Two component UV-curing adhesive (OptiBond, Kerr) was applied to the skull surface and UV cured for 30 – 60 sec at full intensity (Superlite 1300, M+W Dental). A thin layer of metabond was evenly applied to the adhesive surface. A custom-made head-fixation implant (developed by Nelson Totah, Masataka Watanabe, Johannes Boldt and licensed to 3DNeuro, Germany) was attached to the skull via UV-curing cement (Tetric EvoFlow). This cement was UV cured for 60 sec at full light intensity. A craniotomy was made over the cerebellum for a ground wire. The ground wire (silver wire, 0.25 mm diameter, polyimide insulated, Advent Research Materials, AG547509) was stripped of insulation at the tip and flattened using a 3 Ton pneumatic press. The flattened wire was inserted into the craniotomy under the skull. The craniotomy was filled with duragel followed by a biocompatible silicone elastomer (KWIK-SIL, World Precision Instruments). The electrode interface board and wires were packed inside the head chamber and the chamber was filled with 2-component dental cement (Paladur). Rats recovered for at least 5 days after surgery. Carprofen (5.0 mg/kg, s.c.) was administered every 24 hours for 3 days. Rehydrating and easily consumable food was provided in the home cage of the rat (DietGel Recovery).

### Handling and controlled access to water

After recovery, rats were handled and habituated to the setup. Habituation occurred inside the room with the head-fixation apparatus. First, the head-post of the rat was held lightly by hand and the rat was allowed to walk on the treadmill. Daily water intake was controlled to range from 8 to 12 mL, through a combination of water received during the behavioral task and additional water provided in the home cage after the task. Controlled access lasted for 5 consecutive days and was followed by 2 days of ad libitum access.

### Head-fixation and behavioral task apparatus

The head-fixation apparatus and treadmill were inside a 2m x 2m x 2m Faraday cage with a window containing electromagnetic shielding through which the visual stimuli were presented. The rat was head-fixed on a cylindrical, non-motorized treadmill that freely rotates forward and backward. A reward port was placed in front of the mouth of the rat, and 10% sucrose water was delivered via a TTL pulse-controlled pump. Visual stimuli were displayed on a computer screen covering the entire visual field of the rat (60 fps). A rotary encoder was used to measure the angular position of the treadmill. The signal was sampled at 32 kHz (Neuralynx Digital Lynx SX). Velocity (in 5 msec bins) was calculated offline in MATLAB. Videos for facial expression analysis were recorded at 60 fps (Allied Vision Manta G-040B). The camera was facing the right side of the face. Its position was fixed relative to the head across sessions and rats. The camera emitted TTL pulses time-locked to each video frame, and these TTL pulses were recorded in the data acquisition system.

### Behavioral task training

Rats received 40 uL of 10% sucrose in water for small forward body movements. The distance threshold for triggering reward was gradually increased to train the rat to make steps and eventually, to steadily walk or run. Once the rat was licking and walking simultaneously, we trained the rat to make an instrumental response to a Go visual stimulus. Visual stimuli were black and white drifting gratings. During the first step of training, the stimulus was presented for 15 sec. Upon each distance threshold crossing during the stimulus presentation, the rat received a reward delivery. Distance threshold was the same for all rats and sessions. The inter-trial interval was drawn randomly from a distribution from 2 to 3 sec in steps of 0.05 sec. The distribution had a flat hazard rate. After 1 to 2 sessions, rats were continuously walking during the stimulus (and sometimes during the ITI). The number of trials varied and the task was stopped when the rat stopped consuming reward or stopped responding.

The next step of training required the rat to learn to remain immobile prior to stimulus onset. The ITI was decreased to vary between 1 and 2 sec. Running was not permitted in the 0.5 sec before stimulus onset, and resulted in a 0.5 sec time-out and the ITI was restarted. Once rats began to reduce running during the ITI, we reduced stimulus duration in small steps (from 10 sec to 5 sec, to 2.5 sec). Reward was increased to 60 (or 80) uL per threshold crossing for the 5 (or 2.5) sec duration stimulus. Once rats did not respond in the 0.5 sec before stimulus onset at the 2.5 sec stimulus duration level of training, the stimulus duration was further decreased to its final duration of either 1.5 sec or 0.75 sec (all male rats and one female rat were tested with a 0.75 sec stimulus duration, while three other female rats were tested with a 1.5 sec duration stimulus. At the testing stage, the ITI range was changed to 0.5 to 2 sec. At this stage, threshold crossing resulted in 3 pulses of 80 uL reward with 500 msec between pulses. A session was typically 400 to 600 trials (approximately 35 min session duration). Training occurred in one session per day.

In the final step of training, the NoGo stimulus was introduced. Go and NoGo stimuli had at least 70 degrees difference in their orientation. An equal number of Go and NoGo stimuli were presented across 400 to 600 trials in in pseudorandom order (with the limitation that maximally 2 NoGo stimulus trials could occur consecutively due to the difficulty rats have withholding running for multiple NoGo trials and interceding it is). If the rat responded (i.e., crossed distance threshold) in a 1.75 sec window after Go stimulus onset, 3 pulses of 120 uL reward were given separated by 500 msec. The Go stimulus was extinguished upon distance threshold crossing. Responding to the NoGo stimulus also led to extinguishment of the stimulus and a 60 dB, 200 msec duration burst of brown noise followed by a 2 sec time-out. During the ITI and during time-out, a uniform grey screen was presented. Once performance (Hits relative to False Alarms) was above 85% and omission rate was less than 10%, training was complete. Training has been described in detail in our prior work^11,13^.

In order to encourage the rat to commit NMs, we used Go and NoGo stimuli that were difficult to discriminate leading to stimulus confusion and reduced task performance. We targeted, for each rat, the stimulus difference that would produce around 70% correct discrimination. Once training in the Go/NoGo task was completed, rats were run on a psychophysical staircase procedure in which the NoGo stimulus was gradually shifted closer to the Go stimulus by 0.5 degrees in a 2-down / 1-up procedure. The Go stimulus remained the same throughout the experiment. Once the NoGo stimulus converged to a stable level (generating approximately 70% correct performance), this new NoGo stimulus was presented to the rat for around 10 sessions.

### Electrocardiogram data acquisition and R peaks extraction

Signals were recorded at 32 kHz (Neuralynx, Digital Lynx SX) with a bandpass filter of 30 to 1000 Hz. R peaks were detected from the 50 to 150 Hz bandpass filtered signal using a toolbox^38^. Some sessions were bandpass filtered in a narrower to aid in R peak detection. Other parameters such as power, threshold, smoothing and QRS wave window were variable from session-to-session.

### Root mean square of successive differences between R peaks (RMSSD) calculation

Once the R peaks were extracted, we calculated the RMSSD. RMSSD was analyzed as a continuous measure by calculating it in 500 msec sliding windows (100 msec steps). RMSSD was calculated as:

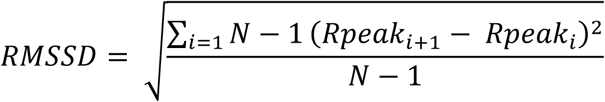

### Facial expression analysis

This analysis was adapted from prior work^10^. Frames of interest, around NM, were taken from the facial videos. All frames were cropped in order to remove most of the space that does not include the rat’s face (cropping coordinates were the same for all rats). Next, frames were converted to black and white and then normalized using power law transformation. Each frame’s main features were described using a HOG (**Supplementary Figure 1A, B**). HOGs for each frame were calculated using the following parameters: 8 histogram orientation bins, using a cell size of [16 16] and 1 cell per block. The cell size was tailored to capture the complexity of the image. For instance, a simple image like a box can be described with a bigger cell size than a cathedral (**Supplementary Figure 1C, D**).

We calculated the pairwise correlations between HOGs T1 NMs and T2 or T3 NMs, using the MATLAB function, corr (**Supplementary Figure 2**). We z-scored normalized the correlations to a baseline epoch (-1 to 0.8 sec before peak velocity for NMs and -0.8 to -0.7 sec before peak velocity for running bouts during ITI) on a trial-wise basis. Pairwise correlations were averaged within rat and then across rats.

### Similarity heatmap between HOGs

All T3 NMs were used to compute the heatmap. On each trial, we took the HOG from the frame that had the lowest correlation with CR face around peak velocity (-0.05 to 0.1 sec around peak velocity). Next we calculated the difference between the T3 NM HOG and the CR HOG. We averaged trials on a session basis, then within rat and across rats, for female and male rats separately. We set all non-face parts of the frame to 0 (this includes the reward port, part of the head chamber and the black space above the ears and nose and below the face) when plotting the result in order to highlight the differences across the face.

## Acknowledgements

This work was funded by a Sigrid Juselius grant (NKT) and the Helsinki Institute of Life Sciences (HiLIFE) at the University of Helsinki (NT).

---

**Supplementary Figure 1.**
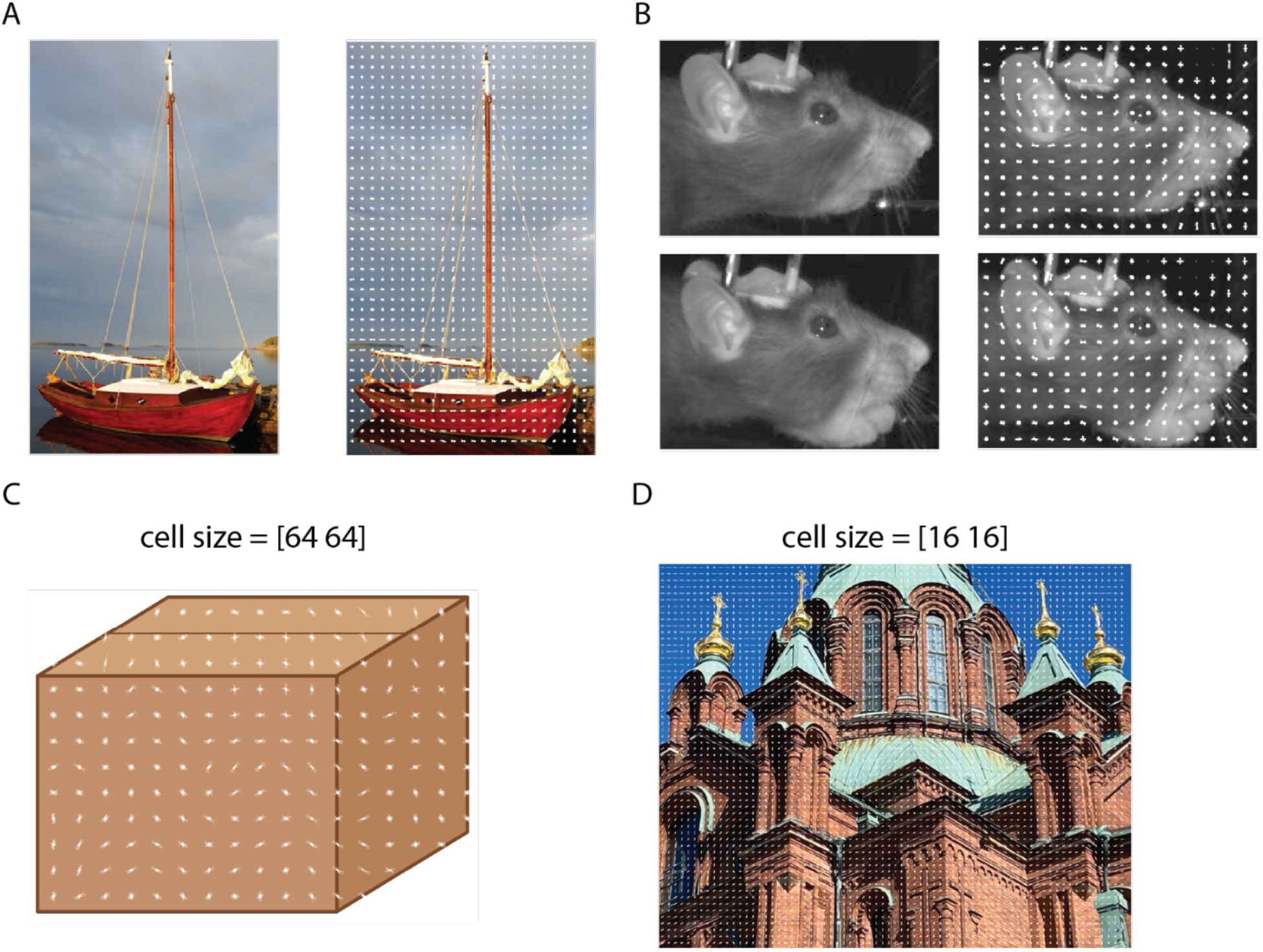
HOG method examples to show how the method depicts picture features and how the pictures influence the parameters of the HOG. Panels **(A)** and **(B)** show, respectively, a picture of a boat and an example frame of a rat’s face together with the HOGs that have been calculated for these images. Each HOG vector represents the changes in contrast, brightness and edges in each pixel. Panels **(C)** and **(D)** demonstrate how the picture’s content will determine how big the cell size will be in order to capture all the features in the picture. A simple box **(C)** requires a higher number of cell size, compared to a picture of a cathedral **(D)**.

**Supplementary Figure 2.**
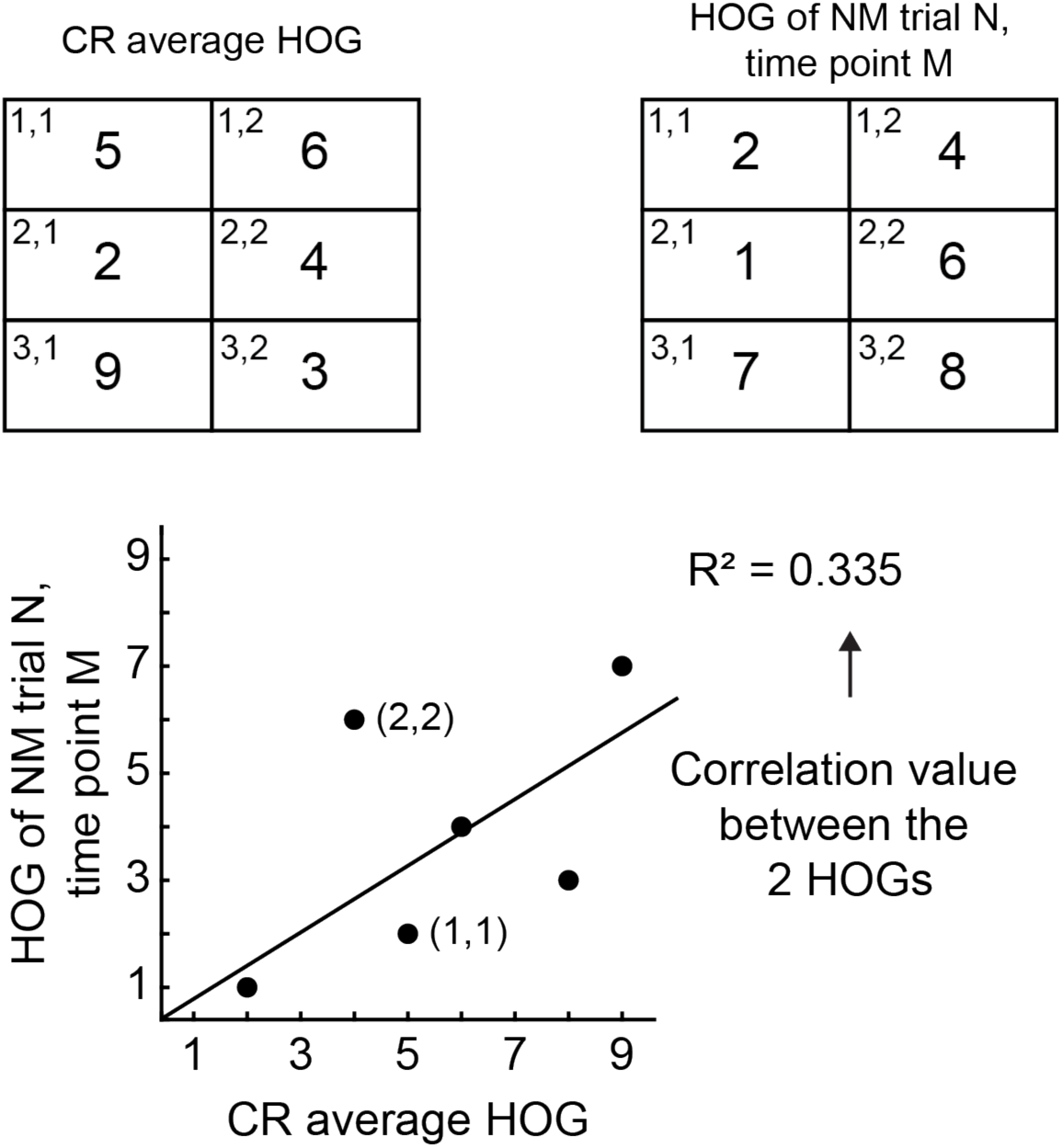
An illustration of how the correlation coefficient is calculated between 2 HOGs. In this hypothetical example, the image has 6 pixels with corresponding coordinates (e.g., “1,1”). The values for each pixel are the result of the HOG analysis. For each time point on each NM trial, the pixel-wise HOGs are compared to the pixel-wise HOGs that are the CR trial- and time-averaged. The pixel-wise correlation is calculated as an R^2^ value. This is repeated for each time point of each NM trial. These values are then z-scored.

**Supplementary Figure 3.**
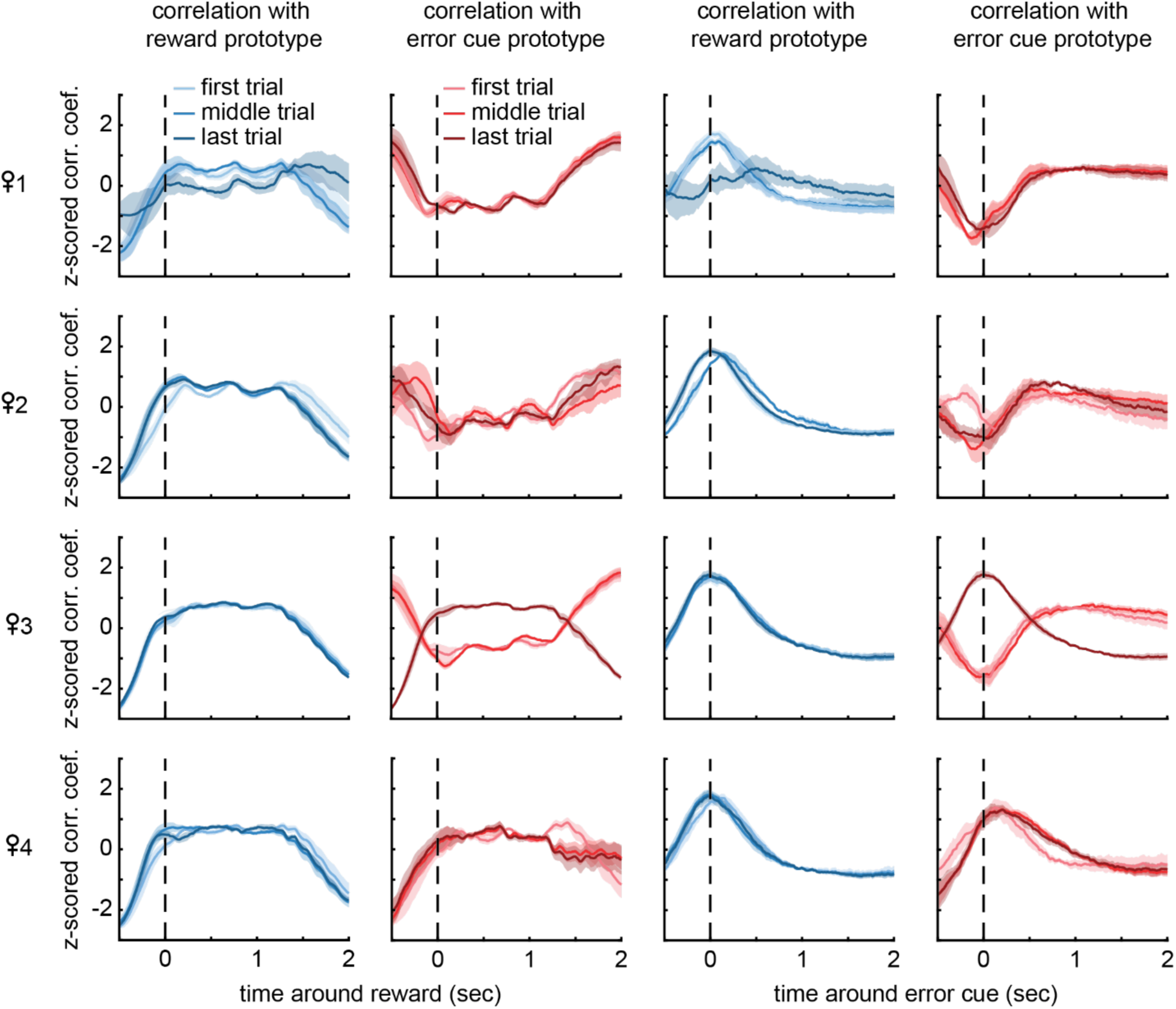
The dependence of correlation coefficients on session time and individual female rat. We calculated z-scored correlation coefficients between facial expression prototypes and task outcomes (reward or error cue) for using each of the four female rats to generate the prototype face (rows of the figure). For each rat, we calculated the correlation using the first trial, session mid-point trial, or final trial. The shading illustrates standard error across sessions for each rat.

**Supplementary Figure 4.**
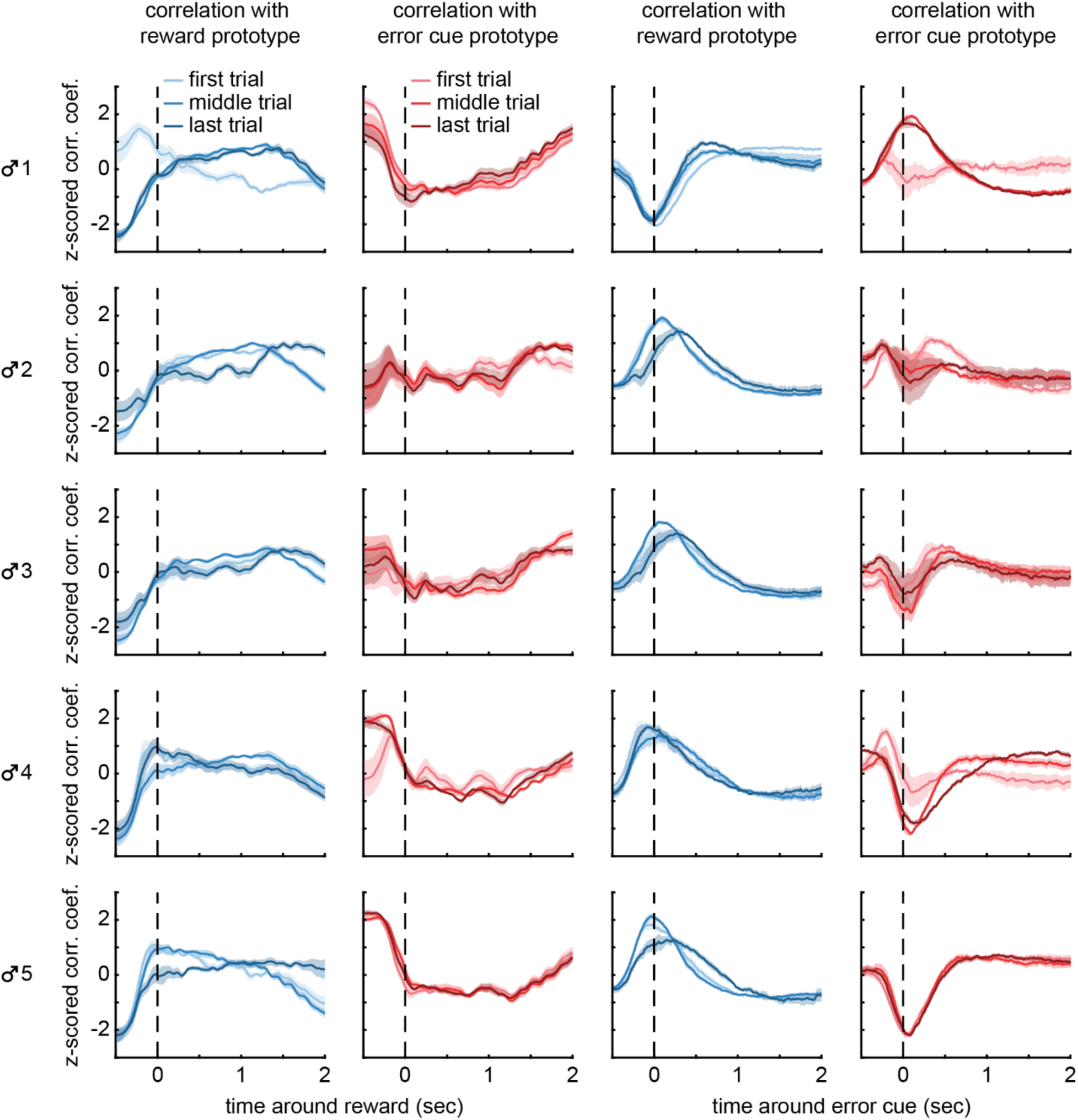
The dependence of correlation coefficients on session time and individual male rat. We calculated z-scored correlation coefficients between facial expression prototypes and task outcomes (reward or error cue) for using each of the five male rats to generate the prototype face (rows of the figure). For each rat, we calculated the correlation using the first trial, session mid-point trial, or final trial. The shading illustrates standard error across sessions for each rat.

